# Rhamnose-mediated modulation of hair follicle growth: insights from chemical genomics

**DOI:** 10.1101/2025.01.26.634903

**Authors:** Xiu Li, Ying Liu, Lu Li, Wenjie Jiang, Shize Ma, Rui Cao, Yi Zhang, Ran Xiao, Dong Wang, Zhigang Yang

**Author notes:** Corresponding authors: Ran Xiao, Dong Wang, and Zhigang Yang., and. These authors contributed equally to this work (Xiu Li, Ying Liu).

## Abstract

Pursuing compounds capable of stimulating hair follicle growth or regeneration presents a significant challenge in dermatological research. Dermal papilla (DP) cells, which serve as a dynamic niche for hair follicle stem cells, are often used to identify hair growth-promoting agents as a screening model. Here, we integrated human scalp single-cell RNA sequencing data with human DP cells (hDPCs) gene expression profiles to develop a gene panel for assessing hair follicle inductive potential, then applied a high-throughput sequencing-based high-throughput screening (HTS^2^) method to quantify the gene expressions of hDPCs perturbed by over 2000 small molecules. Notably, rhamnose was identified as a compound that restored the expression of hair-inducing genes in hDPCs. More importantly, the topical application of rhamnose accelerated hair follicle regeneration by promoting hair cycle into the anagen phase. Transcriptome and central carbon metabolism analyses revealed that rhamnose induced glucose metabolism remodeling favoring glycolysis in mice. A similar phenotype was also observed in rhamnose-treated hDPCs, as demonstrated by glycolysis and mitochondrial stress tests. The present study introduces a novel methodology for evaluating hair-inducing agents and provides valuable insights into the cellular and molecular mechanisms underlying the hair growth-promoting effects of rhamnose. These findings highlight the potential for the therapeutic application of rhamnose in the treatment of alopecia.

## 1 Introduction

Hair loss is a prevalent dermatological affliction that burdens affected individuals psychologically and emotionally. Thus far, clinically effective and validated interventions for alopecia are mainly confined to the utilization of topical minoxidil and oral finasteride (Koralewicz and Szatkowska, 2024). Given the limitation of treatment efficacy and medication-related side effects, searching for novel, more efficacious, and safer therapeutic options is ongoing, with a growing interest in identifying compounds that can stimulate hair follicle regeneration.

The dermal papilla (DP) is a specialized mesenchymal component at the base of the hair follicle, helping to maintain hair follicle structure and regulate its development and cyclic growth by serving as a dynamic niche for a heterogeneous population of multipotent stem cells (Driskell et al., 2011). Previous studies have shown that DP possesses the ability to induce *de novo* hair neogenesis of glabrous skin in hair follicle reconstruction assays, and the quality and quantity of DP cells determine the hair follicle types and size (Andl et al., 2023). DP is also a target for various damaging factors in follicle-related diseases. For instance, in androgenetic alopecia, androgens interact with the DP and alter their secretion of regulatory paracrine factors that govern the proliferation and differentiation of hair follicle stem cells (Oh et al., 2020).

Additionally, human DP cells (hDPCs) are frequently used as a reliable *in vitro* model for evaluating the effects of hair growth-modulating agents at both cellular and molecular levels (Madaan et al., 2018). The one-target, one-drug paradigm has long dominated drug discovery over the past decades. However, compared to drugs that target a single pathway, those that target multiple pathways can potentially overcome the limitations associated with single-target drugs, leading to more effective and safer candidates for disease management. High-throughput sequencing-based high-throughput screening (HTS^2^), which integrates RASL (RNA-mediated oligonucleotide Annealing, Selection, and Ligation) with high-throughput sequencing technology, enables the simultaneous quantitative detection of the expression of hundreds to thousands of genes under various conditions, and has been successfully utilized in various applications, from anti-cancer drug discovery to predicting drug sensitivity and elucidating the underlying mechanisms (Li et al., 2012). Thus, deciphering the gene expression patterns underlying the hair-inductive potential of DP cells, and systematically studying their molecular responses to various conditions through HTS^2^ might be an appropriate approach for hair follicle-promoting drug discovery.

Herein, by integrating public human scalp single-cell transcriptomic datasets and bulk gene expression profiles of DP cultured at different conditions, we first developed a panel to evaluate the hair follicle inductive potential of DP cells. Then, we took hDPCs as an *in vitro* model and applied HTS^2^ to screen more than 2000 compounds for potential therapeutic reagents of hair loss. Among the candidates, rhamnose was identified to restore the expression of hair-inducing related genes in hDPCs and to induce the pre-entry of hair follicles into the anagen in mice. Moreover, transcriptome and central carbon metabolism analyses revealed a compatible metabolic remodeling phenotype in the skin of rhamnose-treated mice, characterized by a significant upregulation of genes encoding key glycolytic enzymes and an increase of intermediate metabolites. Similarly, glycolysis and mitochondrial stress tests indicated that rhamnose induces a metabolic shift from mitochondrial respiration to glycolysis in hDPCs. In general, our research introduces a new methodology to evaluate hair-inducing agents and offers a comprehensive understanding of the cellular and molecular mechanisms underlying the hair growth-promoting effects of rhamnose, which may have important implications for the development of novel therapeutic interventions for hair loss.

## 2 Materials and methods

### 2.1 Single-cell data analysis

The single-cell expression data from GSE193269 and GSE129218 were obtained from the Gene Expression Omnibus (http://www.ncbi.nlm.nih.gov/geo/) database. R package Seurat was applied to analyze the single-cell dataset, including quality control, data filtering and normalization, principal component analysis, Uniform Manifold Approximation and Projection (UMAP) clustering, and data integration (Hao et al., 2021). For doublets, we leveraged the R package DoubletFinder (version 2.0.3) to perform sample-specific doublet inference and subsequent removal (McGinnis et al., 2019). By setting the expected doublet rate to 5% and utilizing default parameters, we minimized the impact of any potential doublets on downstream analyses.

### 2.2 Cell isolation and culture

Human normal scalp tissue samples were obtained from patients undergoing plastic surgery procedures, and the collection was approved by the Ethical Committee of Plastic Surgery Hospital Peking Union Medical College (PSH, PUMC). Written informed consent was obtained from all subjects. The scalp samples were initially cut into thin strips (1 x 0.2cm) using a blade and the deep subcutaneous adipose tissue was removed carefully. The strips were then cut apart at the boundary of the dermis-subcutis, and the dermis-subcutis tissues were minced into 2-3 mm^3^ and digested in 0.2% type I collagenase (#C0130; Sigma-Aldrich, Darmstadt, Germany) solution at 37°C for 3.5 hours. Thereafter, the dermal papillae were picked out under a microscope and cultured in DMEM (#10-013-CV; CORNING, Arizona, United States) containing 10% fetal bovine serum (#A5669801; ThermoFisher, Massachusetts, United States) and 1% penicillin-streptomycin (#15140148; ThermoFisher, Massachusetts, United States). For dermal fibroblasts, the cut scalp strips with epidermis were immersed in 0.25% dispase II (#04942078001; Roche, Basel, Switzerland) solution overnight at 4°C. After removing the epidermis, the dermal tissues were digested and the fibroblasts were isolated and cultured. All cells were maintained in a standard humidified incubator maintained at 5% CO_2_ at 37 °C.

### 2.3 Alkaline phosphatase staining

Cells were fixed with 4% paraformaldehyde in phosphate-buffered saline (PBS) for 10 min and then incubated with an alkaline phosphatase staining solution (#SCR004; Sigma-Aldrich, Darmstadt, Germany) at 37°C for 10–15 min. The reaction was stopped by washing cells with PBS, and the images were taken under Microscope (Leica, Wetzlar, Germany).

### 2.4 RNA-seq and data analysis

RNA extraction was performed using TRIzol reagent, followed by mRNA purification using poly-T oligo-attached magnetic beads. Subsequently, sequencing libraries were prepared using the NEBNext^®^ UltraTM RNA Library Prep Kit for Illumina^®^ (NEB, Massachusetts, United States) according to the manufacturer’s protocols. Index codes were introduced to assign sequences to respective samples. The index-coded samples were then clustered using the TruSeq PE Cluster Kit v3-cBot-HS (Illumina, Massachusetts, United States) on the cBot Cluster Generation System. Finally, library preparations were subjected to Illumina NovaSeq sequencing, generating 150 bp paired-end reads.

After quality control, the clean reads were mapped to the reference genome (mm10 for mice samples, hg19 for human samples) using HISAT2. featureCounts v1.5.0-p3 was used to count the read numbers mapped to each gene. Then, the FPKM of each gene was calculated based on the length of the gene and the read count mapped to this gene. Differentially expressed genes (DEGs) were identified based on an adjusted *p*-value threshold of <0.05 using the DESeq2 package (Love et al., 2014). Functional annotation and enrichment analyses were performed using the R package clusterProfiler 4.0 (Wu et al., 2021) and then visualized using the ggplot2 package (Ginestet, 2011). Hierarchical clustering was generated using the pheatmap package (https://cran.r-project.org/web/packages/pheatmap/index.html). A scissor package was used to integrate bulk and single-cell data to identify cell subpopulations affected by rhamnose (Sun et al., 2022). For the dichotomous variable phenotype, Scissor returned Scissor+ cells that stand for ones responding to rhamnose, while Scissor− cells stand for ones associated with the control.

### 2.5 Drug screening and data processing

Approximately 3,000 hDPCs were seeded per well in 384-well plates. After 24 hours, the cells were treated with small molecules from the screening library at a concentration of 1.25 μmol/L for an additional 24 hours. Probes were designed and tested for each gene. HTS^2^ assay was then performed to quantify the mRNA level of the target genes, following the protocol described (Li et al., 2012). Cell lysis was performed using GentLys buffer (Nanopure, China). Sequencing reads were aligned to the probe sequences, allowing up to three mismatches, and the read counts were normalized to the expression levels of 8 stable housekeeping genes. The Pearson correlation coefficients for the normalized transcriptional data were calculated using R software, based on treatments with 16 DMSO replicates. Notably, correlation coefficients of >0.9 demonstrated that the HTS^2^ assay results were reliable and reproducible. For each compound treatment, fold changes in gene expression were calculated by comparing the normalized expression of treated samples to the average normalized expression of DMSO-treated controls within the same 384-well plate. Genes with |log_2_(fold change)| >1.5 and p-value <0.05 were classified as differentially expressed genes (DEGs) using the DESeq2 package (Love et al., 2014). The heatmap of DEGs was generated using the pheatmap and ggplot2 package (Ginestet, 2011).

### 2.6 Anagen induction assay in C57BL/6 mice

All animal experiments were conducted following the guidelines and prior approval of the Animal Care and Use Committee of the PSH, PUMC. 7-week-old female C57BL/6 mice (SPF Biotechnology Co., Ltd, Beijing, China) were housed under SPF conditions operated by the Center of Experimental Animals in the PSH, PUMC. After the back skin of mice in the early telogen phase was shaved with a clipper, Rhamnose (#R3875; Sigma-Aldrich, Darmstadt, Germany) at a concentration of 10 mM and vehicle (0.1% sodium hyaluronate, Yeasen Biotechnology, Shanghai, China) were applied topically to the back of mice daily for 3 weeks. Three independent replicate experiments were performed. The relative areas of skin with darkening color or neonatal hair coverage were measured using the Image J software. Hematoxylin and eosin (HE) staining and immunofluorescence staining were applied for histological analysis of hair follicle growth.

### 2.7 Immunofluorescence

Immunofluorescence staining for Krt15 (#ab52816; Abcam, Cambridge, United Kingdom), Ki67 (#ab16667; Abcam, Cambridge, United Kingdom), PDK1 (#ab202468; Abcam, Cambridge, United Kingdom), and PDH E1 Alpha (#18068-1-AP; Proteintech, Chicago, United States) were performed on 5-μm paraffin sections of mice skin tissues as described in the protocol of the manufacturer’s kit (#abs50029; absin, Shanghai, China). Briefly, slices were stained with primary and TSA secondary antibodies sequentially. Amplification of signals was then carried out, followed by treatment with antibody eluent. Subsequently, the slices were stained with the second antibody using a similar procedure. Fluorescent images were acquired using confocal microscopy (Leica, Wetzlar, Germany).

### 2.8 Metabolomics analysis

After shaving the dorsal hair of mice in the early stages of the telogen phase, 10 mM rhamnose and solvent were applied daily to the dorsal skin for 7 consecutive days. A 100 mg sample of dorsal skin was immediately flash-frozen in liquid nitrogen. Each sample was precisely weighed and transferred to an Eppendorf tube. After the addition of two small steel beads and 500 μL of precooled methanol/water (3:1, v/v; -40°C), samples were vortexed for 30 seconds, homogenized at 40 Hz for 4 minutes, and sonicated in an ice-water bath for 5 minutes. This homogenization and sonication cycle was repeated three times. Samples were then incubated at -40°C for 1 hour and centrifuged at 13800 (× g) at 4°C for 15 minutes. The supernatants (400 μL) were collected and evaporated to dryness by spin. The residue was reconstituted in 200 μL of water, vortexed, filtered, and transferred to injection vials for high-performance Ion chromatography-tandem mass spectrometry (HPIC-MS/MS) analysis. Standard solution preparation, instrument parameter setting, and data analysis methods, were performed as the protocol described (Yang et al., 2023).

### 2.9 Glycolysis stress and Mitochondrial stress test

hDPCs of passage one were seeded at a density of 10,000 cells per well in XF24 cell culture plates and then clustered with DMEM containing 10% fetal bovine serum at 37°C. When cells reached approximately 90% confluency, the culture media was removed and cells were washed twice with a pre-warmed XF-base medium. To perform the glycolytic stress test (#103020-100; Agilent, California, United States), the cells were treated with 500 μL of XF-based medium containing 2 mM glutamine followed by one-hour incubation in a non-CO_2_ humidified chamber. In the meantime, the sensor cartridge was hydrated in a non-CO_2_ incubator overnight at 37 °C. Subsequently, hDPCs were subjected to glycolytic function analysis using the Agilent Seahorse Analyzer according to the standard protocol. For the mitochondrial stress test (#103015-100; Agilent, California, United States), 500 μL of XF-base medium containing 1 mM pyruvate, 2 mM L-Gln, and 10 mM glucose was added to the cells before loading into the Agilent Seahorse Analyzer. Standard measurements were executed following sequential treatment with 1.5 μM oligomycin, 2.0 μM FCCP [Carbonyl cyanide-4 (trifluoromethoxy) phenylhydrazone], and 0.5 μM rotenone/antimycin A. Each assay was run in duplicate. The obtained data were normalized against the total protein content within each well, which was quantified using a BCA Protein Assay kit (#P0012; Beyotime, Beijing, China).

### 2.10 Statistical analysis

Statistical analyses were performed using Prism 8.0 software (GraphPad Software, Inc., United States). All data were presented as the mean ± standard error of the mean (SEM). Unless otherwise indicated, the comparisons between the groups were conducted using Student’s t-test. Statistical significance was defined as *p* < 0.05.

### 2.11 Study approval

Human scalp tissues were obtained from donors during plastic surgery procedures. The Ethical Committee of Plastic Surgery Hospital, CAMS&PUMC approved the collection, and written informed consent was obtained from all subjects. All animal experiments were conducted following the guidelines and prior approval of the Animal Care and Use Committee at the Plastic Surgery Hospital, CAMS&PUMC.

### 2.12 Data availability

The datasets presented in this study can be found in online repositories. The raw sequence data have been deposited in the National Genomics Data Center under BioProject accession number: PRJCA033327.

## 3 Results

### 3.1 HTS^2^ screening identified rhamnose as a potential hair growth-promoting compound

To gain insights into the molecular signature that underlies the hair inductive potential of hDPCs, we first reanalyzed the single-cell transcriptomic datasets of human scalp tissues (Shim et al., 2022) and revealed 19 distinct cell clusters according to their gene expression profiles, and identified 120 marker genes for hDPCs (Figure 1A&B, Figure S1A&B). We also focused on genes reprogrammed during the 3D culture process, which can prevent hDPCs from losing their hair induction ability after *in vitro* expansion (Higgins et al., 2013). By integrating these two datasets, we developed a gene panel to assess the hair-inducing potential of DP cells cultured *in vitro* (Figure 1C, Figure S1C). The panel consists of 35 hDPC signature genes with an increase or restoration in expression following a transition from 2D to 3D spheroid culture, 14 genes with a decreased expression upon hDPCs restoring their hair-inducing capacity, and 2 genes stable across both culture conditions (Figure 1C).

**Figure 1.**
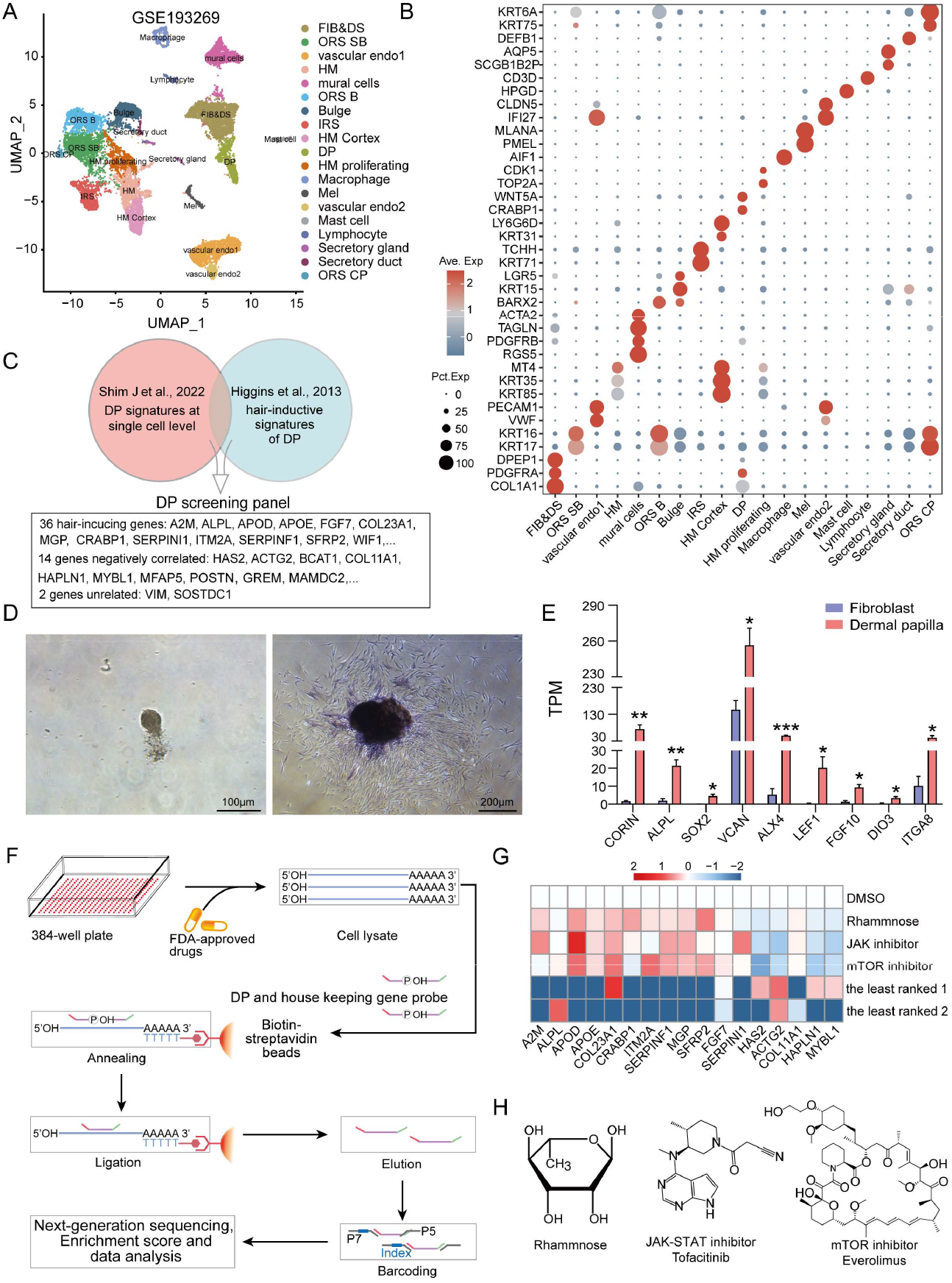
HTS^2^ screening identified compounds with potential hair growth-promoting effects. (A) UMAP visualization of cell clusters within human scalp tissue. (B) Dotplot displaying distinct marker genes for different cell clusters. (C) Schematic diagram illustrating the panel containing 52 genes from two datasets. (D) The morphology of freshly isolated dermal papilla from the digested tissue (left) and ALP staining of cultured hDPCs (right). (E) Bulk RNA-seq data showing the differential expressions of DP signature genes in hDPCs and fibroblasts. (F) Schematic illustration of the experimental procedure of HTS^2^. (G) Heatmap representing the expression of partial panel genes in hDPCs treated with the top and least-ranked compounds. (H) The selected chemical structures are shown. TPM, Transcripts Per Million. *P < 0.05, **P < 0.01, ***P < 0.001. FIB, Fibroblast; DS, Dermal sheath; ORS SB, Suprabasal outer root sheath; HM, Hair matrix; ORS B, Basal ORS; IRS, Inner root sheath; Mel, Melanocyte; ORS CP, ORS companion layer.

hDPCs were then isolated from scalp tissue and cultured in an adherent manner. Consistent with previous reports (Wei et al., 2021), the cells displayed positive staining of alkaline phosphatase and much higher expression levels of *CORIN, ALPL, SOX2, VCAN, ALX4, LEF1, FGF10, DIO3*, and *ITGA8* than those in fibroblasts, suggesting that the hDPCs were successfully isolated with high purity (Figure 1D&E, Figure S1D). hDPCs were then treated with a library of 2145 small molecules, including FDA-approved drugs, bioactive compounds, and natural products, and the mRNA levels of panel genes were profiled using the HTS^2^ method (Figure 1F). The activity of each compound was evaluated based on its capacity to restore the gene expression levels that correlated with the hair-inducing ability (Figure S1E). Among the positive compounds with the potential to promote the hair-inducing ability of hDPC, two compounds targeting the JAK-STAT and mTOR signaling, promote the expression of *A2M, APOD, APOE, COL23A1, SFRP2, FGF7*, etc., and inhibit *HAS2, ACTG2, COL11A1, HAPLN1*, and *MYBL1*, whereas the potential negative regulator displayed reverse effects on the expression of the aforementioned genes (Figure 1G). Both JAK-STAT and mTOR signaling inhibitors have been reported to stimulate hair growth and initiate the hair cycle transition from telogen to anagen (Chai et al., 2019, Harel et al., 2015), suggesting our approach can effectively identify potential hair-promoting drugs. Of note, we found L-rhamnose (6-deoxy-L-mannose), a natural sugar widespread in plants and bacteria, exhibited similar effects to JAK-STAT and mTOR inhibitors, suggesting L-rhamnose may also be a promising candidate (Figure 1G&H).

### 3.2 Rhamnose induced hair cycle transition from telogen to anagen in mice

To assess the potential effects of rhamnose on hair growth *in vivo*, we topically applied 1 mM and 10 mM rhamnose to the shaved dorsal skin of 7-week-old C57BL/6 mice. These mice were at the second telogen phase that typically lasts for about a month, and were used as a classic model to assess the efficacy of compounds in stimulating hair follicle growth (Geyfman et al., 2015). The hair growth status was recorded during the subsequent observation period. As shown in Figure 2A, some mice showed significant skin pigmentation and hair growth on the dorsal and caudal areas of skin as early as 14 days especially after 10 mM rhamnose treatment, while the controls remained unpigmented. From day 14 to day 21, the number of mice with expanded skin darkening areas or hair growth on their dorsal skin was much higher in the rhamnose-treated group than in the control group (Figure 2A). Since melanogenesis is coupled to anagen initiation in murine, we measured the area of pigmented skin patches and found a significant increase in mice treated with 10 mM rhamnose at all checked time points, including days 14, 18, and 21 (Figure 2B). The same increasing tendency was observed for areas covered by newly formed hairs in rhamnose-treated mice compared with the controls (Figure 2C). Hematoxylin and Eosin (HE) staining of dorsal skin tissues showed that most of the control mice displayed a telogen or early anagen morphology with hair bulbs located in the dermis, whereas the hair follicles of rhamnose-treated mice exhibited classical anagen structure with significantly enlarged bulbs extending into the subcutaneous adipose tissue (Figure 2D). Consistently, immunofluorescence staining revealed that Ki67, a well-established marker for cellular proliferation, exhibited high and widespread expression in the cells of hair bulbs from the rhamnose-treated group, but was scarcely detected in the control group (Figure 2E). These findings strongly suggested that rhamnose significantly promoted the telogen-to-anagen transition of mice hair follicles.

**Figure 2.**
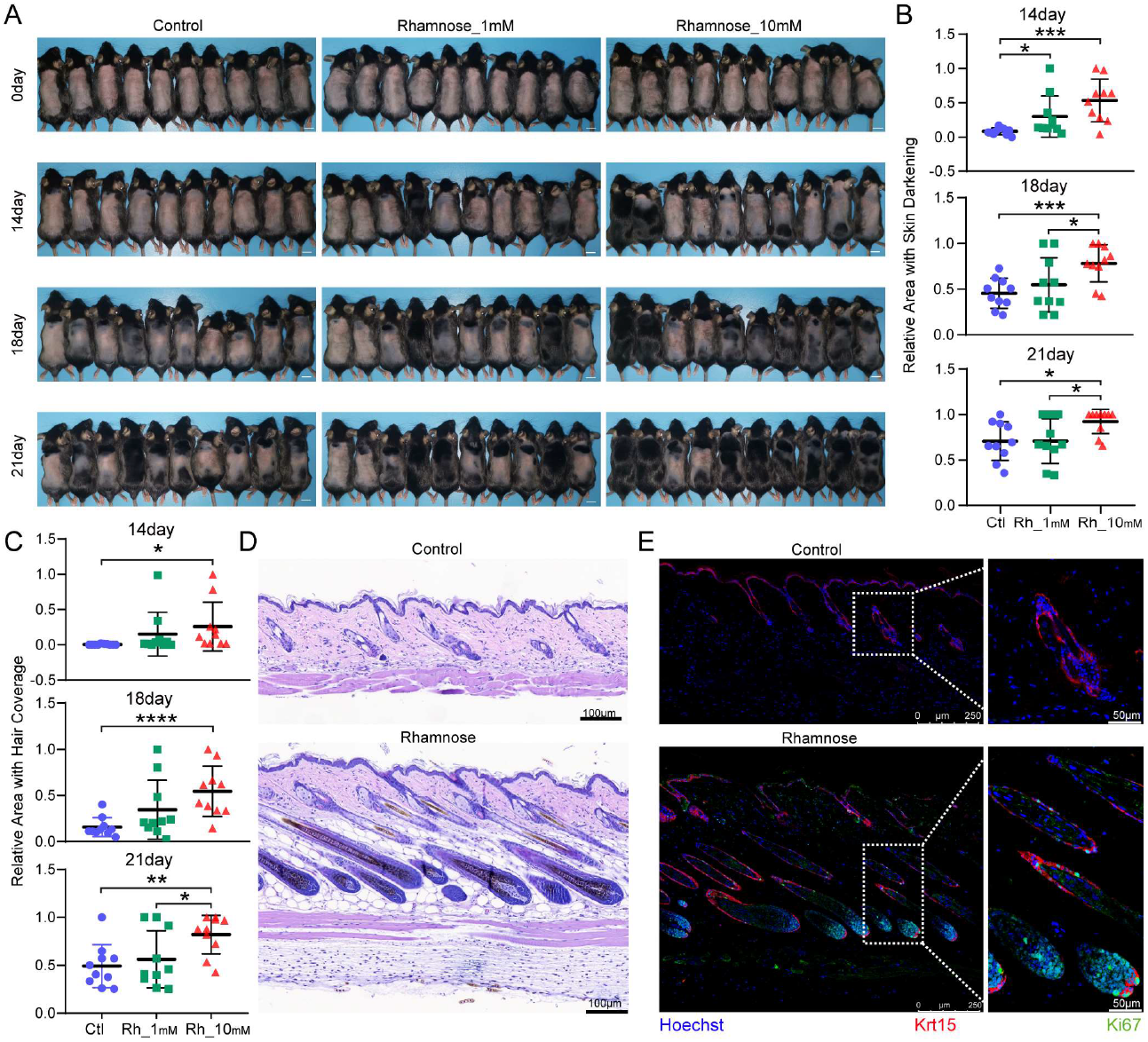
Rhamnose promoted anagen induction in C57BL/6 mice. (A) Mice with shaved dorsal hair were photographed at 0, 14, 18, and 21 days after vehicle and rhamnose treatment. (B, C) Statistic analysis of the relative area of skin pigmentation (B) and hair coverage (C) in the dorsal skin of mice from vehicle and rhamnose-treated mice. (D) Representative Hematoxylin and Eosin (H&E) staining of dorsal skin samples after a 21-day treatment with both vehicle and rhamnose. (E) Immunofluorescent staining showing the expression of keratin 15 and Ki67 in the dorsal skin samples following a 21-day treatment with vehicle and rhamnose. *P < 0.05, **P < 0.01, ***P < 0.001, ****P < 0.0001. Ctl, Control, Rh_1mM, Rhamnose_1mM, RH_10mM, Rhamnose_10mM.

### 3.3 Rhamnose promoted anagen initiation at both cellular and molecular levels

To further investigate the effects of rhamnose on hair follicle growth, skin tissues from pre-treatment, 7-, 10- and 14-day rhamnose or vehicle-treated mice were subjected to transcriptome sequencing analysis. Consistent with the skin phenotype, PCA analysis revealed that rhamnose-treated biological replicates at 14 days formed a distinct cluster and showed less similarity to other groups, suggesting a dramatic change in the hair follicle status after 14 days of treatment with rhamnose (Figure 3A). In the skin of 14-day rhamnose-treated mice, we found 3516 upregulated genes and 4393 downregulated genes, as compared to the pre- and vehicle-treated groups (Figure S2A). The Gene Set Enrichment Analysis (GSEA) demonstrated that the upregulated genes were enriched in biological processes such as cell proliferation and division, epidermis development, hair follicle morphogenesis, and melanin synthesis (Figure S2B). This was further supported by CNET plot highlighting key genes involved in these processes, including Msx2, Cdh3, Wnt5a, Shh, Foxe1, and Foxn1, among others (Figure S2C), and by a heatmap displaying dramatic upregulation of genes associated with the hair follicle epithelium (e.g., *Krt6a, Krt17, Krt71, Krt83, Lgr5*), anagen DP (e.g., *Hhip, Rspo2, Edn3, Scube3*), and melanocytes (e.g., *Tyr, Mlana, Pmel, Gpnmb*) (Figure 3B).

**Figure 3.**
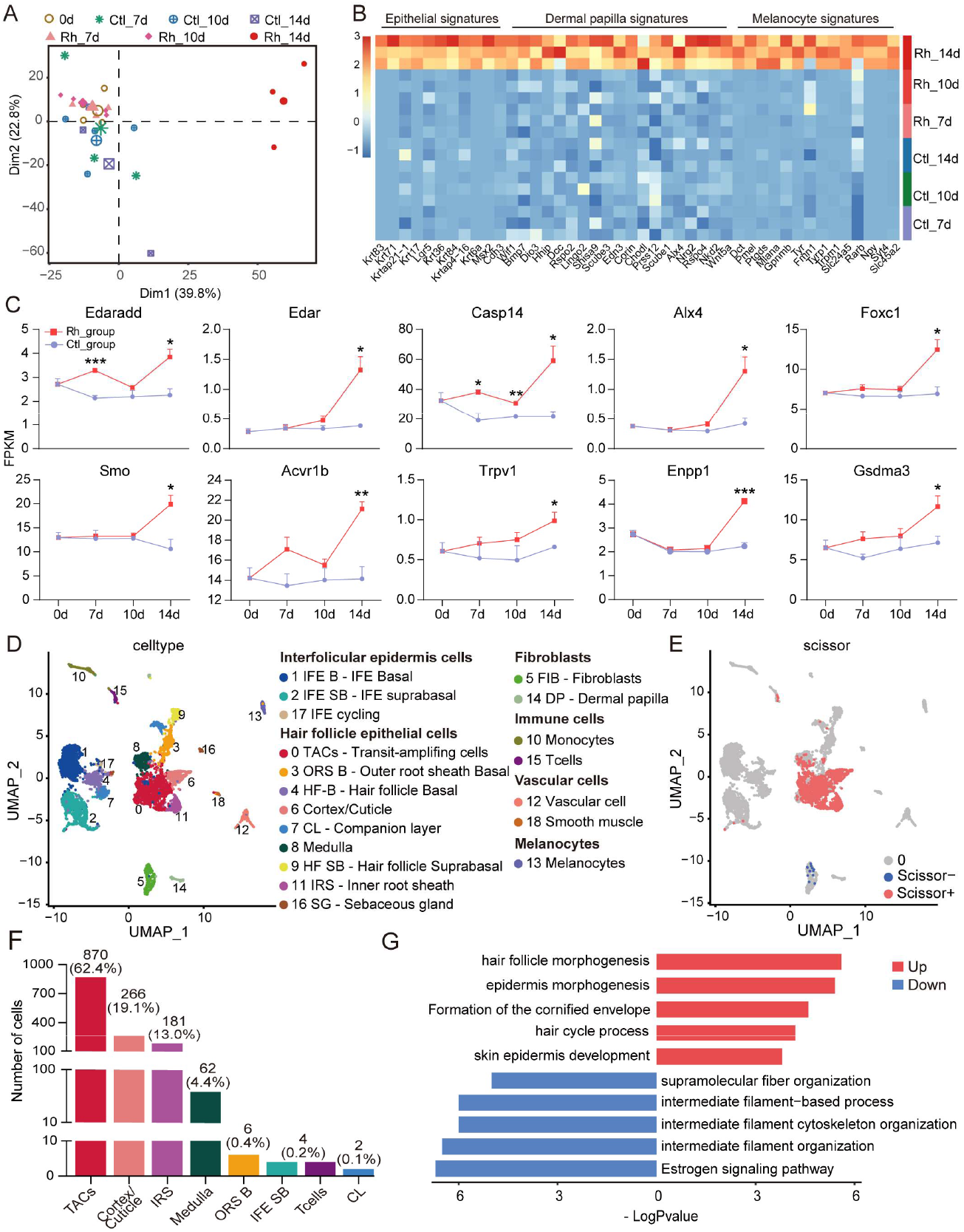
Identification of cell subpopulations and molecular changes associated with rhamnose treatment. (A) PCA plot showing transcriptome data from pre-, vehicle-, and rhamnose-treated groups at 7, 10, and 14 days. (B) Heatmap showing highly expressed genes specific to epithelial cells, melanocytes, and DP in the skin of 14-day rhamnose-treated mice. (C) Expression trends of genes related to hair follicle development and growth in vehicle-, and rhamnose-treated groups across different time points. (D) The UMAP clustering of single cells from female C57BL/6 skin at both anagen and telogen phases. (E) The UMAP distribution of the scissor-selected cells. (F) Bar plot illustrating the detailed distribution of cell types within the scissor-selected cell population. (G) Significantly enriched pathways in Scissor+ cells. Ctl_7d, Control_7day, Ctl_10d, Control_10day, Ctl_14d, Control_14day, Rh_7d, Rhamnose_7day, Rh_10d, Rhamnose_10day, Rh_14d, Rhamnose_14day, Rh_group, Rhamnose_group, Ctl_group, Control_group.

We also evaluated the dynamic expression of key genes related to hair follicle development and growth during the treatment process. We found that rhamnose upregulated the expression of certain genes as early as 7 days into the treatment, such as *Edaradd* and *Casp14*. Edaradd coexpresses and interacts with Edar, acting as an adaptor to link the receptor to downstream signaling pathways. It plays an important role in epithelial cells during the formation of hair follicles and teeth (Gao et al., 2023). Casp14, a non-apoptotic caspase, is expressed in the differentiating and cornifying layers and hair follicles and is involved in epidermal differentiation (Rendl et al., 2002). Other important players in hair follicle development and growth, such as the transcription factors *Alx4, Foxc1*, as well as Hedgehog receptor *Smo*, were significantly upregulated at 14 days of treatment. These genes showed an increasing tendency in the early days of treatment, although the changes were not statistically significant (Figure 3C).

To explore the cellular change induced by rhamnose treatment, we integrated our bulk RNA-seq data with the GSE129218 single-cell dataset and applied the Scissor algorithm to identify specific cell types that changed in rhamnose-treated mouse skin. Scissor is a novel single-cell data analysis approach that uses bulk phenotypes to guide single-cell data analysis, enabling hypothesis-independent identification of clinically and biologically relevant cell subpopulations. The GSE129218 dataset is from full-thickness female C57BL/6 skin during anagen (5w) and telogen (9w). As shown in Figure 3D, we identified 19 cell clusters including interfollicular epidermal cells, hair follicle epithelial cells, fibroblasts, vascular cells, immune cells, melanocytes, and Schwann cells (Figure 3D, Figure S2D). After integrating DEGs of 14-day rhamnose-treated skin tissues, we identified 1395 Scissor+ cells. Remarkably, 99.9% of these cells were from anagen skin, whereas only 12 Scissor-cells were found, mainly present in the telogen phase (Figure 3E, Figure S2E&F). Specifically, Scissor+ cells predominantly distributed in clusters such as TACs, IRS, cortex/cuticle, and Medulla, and exhibited high expression levels of hair follicle development and cycling growth-related genes, such as hair cortex/cuticle marker Krt35, IRS markers *Krt25, Krt27, Krt28*, transcription factor *Lef1*, and hair matrix TACs marker *Dcn* (Figure 3F&G, Figure S2G&H). Overall, these findings indicated rhamnose promotes the expression of key genes involved in hair follicle growth and induces cellular reconstitution during anagen initiation.

### 3.4 Rhamnose elicited the remodeling of genes associated with glycolysis and the TCA cycle in the skin during the early period of treatment

Upon transition from telogen to anagen, the cellular composition and size of HFs change significantly. We reasoned that any transcriptome changes at this phase would be the consequence of anagen phenotype rather than its cause. Therefore, we focused on the transcriptomic data from rhamnose-treated samples at 7 and 10 days, where no obvious phenotypic changes were observed, and we identified significant alterations in gene expression profiles within the skin tissue following treatment of 7 and 10 days. Compared to the control on the same day, rhamnose induced 2150 and 2228 DEGs in 7- or 10-day treatment, respectively (Figure 4A&B). These findings suggested that rhamnose elicits transcriptional changes in the skin at an early stage, prior to the manifestation of discernible phenotypic effects. Enrichment analysis revealed distinct biological processes associated with DEGs at different time points. While DEGs at 14 days were significantly involved in ‘Extracellular matrix organization’, ‘Mitotic Prometaphase’, and ‘Keratinization’, the genes that exhibited changes at 7 days after rhamnose treatment were predominantly engaged in ‘Glycosaminoglycan metabolism’, ‘Metabolism of lipids’, and ‘Glycogen metabolism’. Furthermore, DEGs after 10 days of treatment were notably enriched in ‘Glycolysis’, ‘TCA cycle’, and ‘Respiratory electron transport’ pathways (Figure 4C). The CNET plots further revealed the biological concept related to gene changes at each time point (Figure 4D, Figure S3).

**Figure 4.**
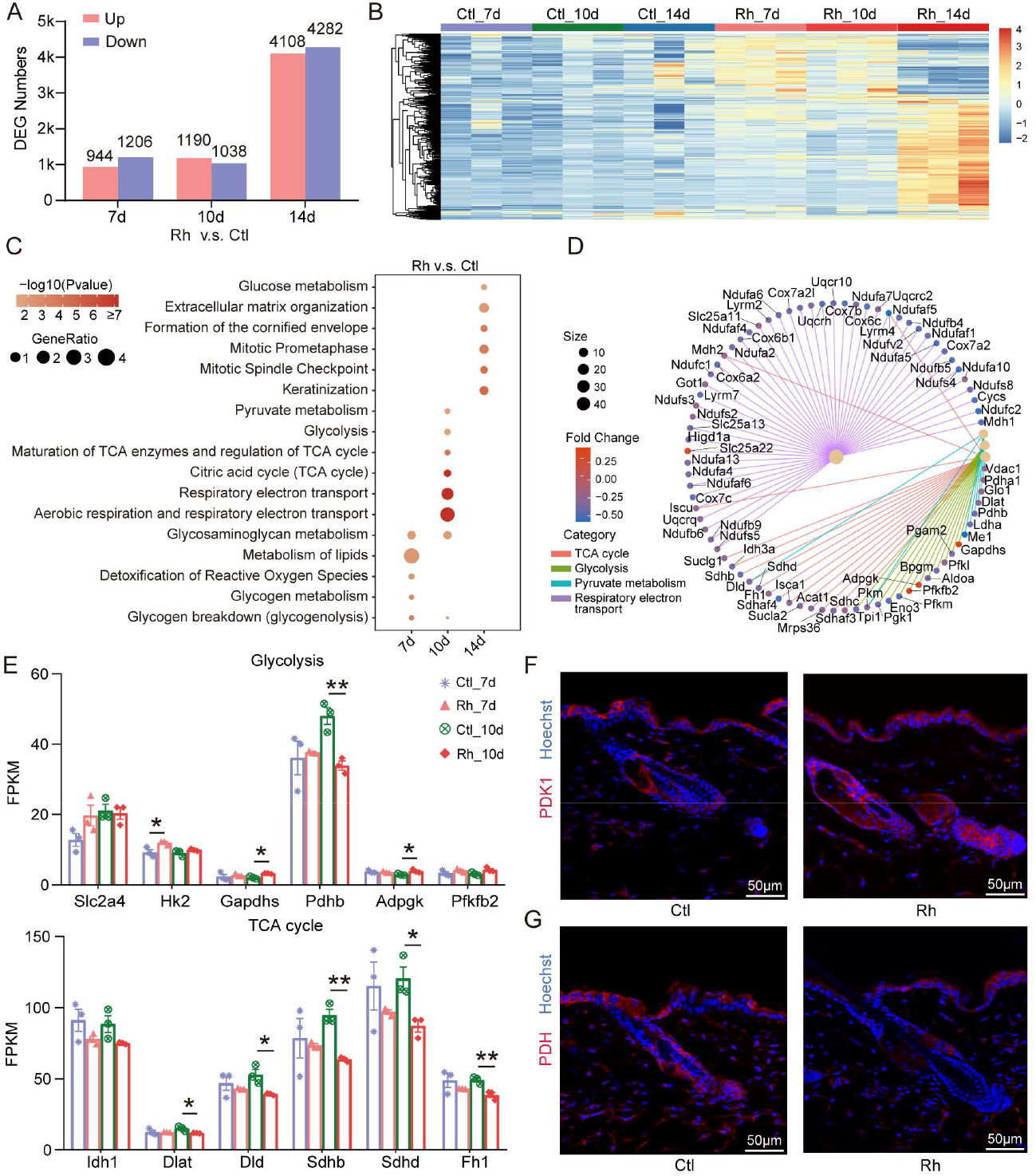
RNA sequencing revealed metabolic changes in mouse skin treated with rhamnose. (A) The numbers of DEGs between rhamnose and control groups at the different days after treatment. (B) Hierarchical clustering heatmap showing different gene expression profiles across samples. (C) Significantly enriched Reactome pathways in rhamnose-treated groups at 7-, 10-, and 14-day. (D) CNET plot of the representative biological processes enriched in the rhamnose-treated 10days group. (E) Expression profiles of genes associated with glycolysis and the TCA cycle across various groups. (F, G) Immunofluorescent staining showing the expression of PDK1 (F) and PDH (G) in the dorsal skin samples following 7 days of treatment with vehicle and rhamnose. *P < 0.05, **P < 0.01. Ctl_7d, Control_7day, Ctl_10d, Control_10day, Rh_7d, Rhamnose_7day, Rh_10d, Rhamnose_10day.

Among these, glycolysis-related genes, including glucose transporter GLUT4 (*Slc2a4*), hexokinase II (*Hk2*), which catalyzes the rate-limiting first obligatory step of glucose metabolism, Glyceraldehyde-3-phosphate dehydrogenase (*Gapdhs*), ADP-specific glucokinase (*Adpgk*), and 6-phosphofructo-2-kinase/fructose-2,6-bisphosphatase 2 (*Pfkfb2*), were significantly upregulated (Figure 4E). In contrast, genes involved in the TCA cycle, such as key enzymes *Idh1, Sdhb, Sdhd*, and *Fh1*, were consistently downregulated following rhamnose treatment (Figure 4E). Meanwhile, components of the pyruvate dehydrogenase complex, such as *Dlat, Dld*, and the β-subunit of the pyruvate dehydrogenase (*Pdhb*), which are located in mitochondria and catalyze the conversion of pyruvate to acetyl-CoA, were also downregulated after rhamnose treatment (Figure 4E). Immunofluorescence staining also verified that PDH was downregulated, whereas pyruvate dehydrogenase kinase 1 (PDK1), which phosphorylates PDH and results in its inactivation, showed a significant increase in expression following rhamnose treatment (Figure 4F&G). These coordinated changes in gene expression induced by rhamnose treatment suggested an enhancement of glycolytic activity and a reduction in the TCA cycle activity.

### 3.5 Rhamnose increased major intermediate metabolites of glycolysis

Central carbon metabolism (CCM) is the fundamental metabolic process that maintains cellular growth and provides precursors for biosynthesis. To verify the potential impact of rhamnose on the metabolic phenotype of the skin, we conducted comprehensive metabolite profiling analysis on mouse skin samples following a 7-day regimen with either 10 mM rhamnose or vehicle treatment. We reasoned metabolic alterations occurring at such an early stage would likely be more easily understood as the cause, rather than the consequences of the anagen phenotype. This analysis enabled the absolute quantification of 56 metabolites, which were categorized into several key metabolic pathways: the pentose phosphate pathway, the tricarboxylic acid (TCA) cycle, and the glycolysis and gluconeogenesis pathways (Figure 5A). The PCA score plot revealed a marked separation between control and rhamnose-treated groups, indicating substantial metabolic differences between these conditions (Figure 5B). Specifically, 45 metabolites were identified, and the hierarchical clustering analysis showed that rhamnose significantly altered the glycometabolite profiles in the skin (Figure 5C). Of note, major glycolytic intermediates, such as glucose-6-phosphate, fructose-6-phosphate, fructose-1,6-bisphosphate, dihydroxyacetone phosphate, 3-phosphoglyceric acid, and phosphoenolpyruvic acid, were significantly enriched in the rhamnose-treated group compared to the control, indicating an increase in glycolytic activity (Figure 5D). In contrast to the glycolysis pathway, most of the metabolites involved in the TCA cycle were not changed after rhamnose treatment, though cis-aconitic acid and isocitric acid showed a slight increase (Figure S4A). For the pentose phosphate pathway, we observed an upregulation of 6-phosphogluconic acid and erythrose 4-phosphate, concurrent with the downregulation of gluconic acid and ribose 5-phosphate (Figure S4B). Consistent with the transcriptomic results that rhamnose increases the expression of genes encoding key glycolysis enzymes, our metabolomic findings also indicated that rhamnose remodels the glucose metabolism processes in mouse skin to support glycolysis (Figure 5E).

**Figure 5.**
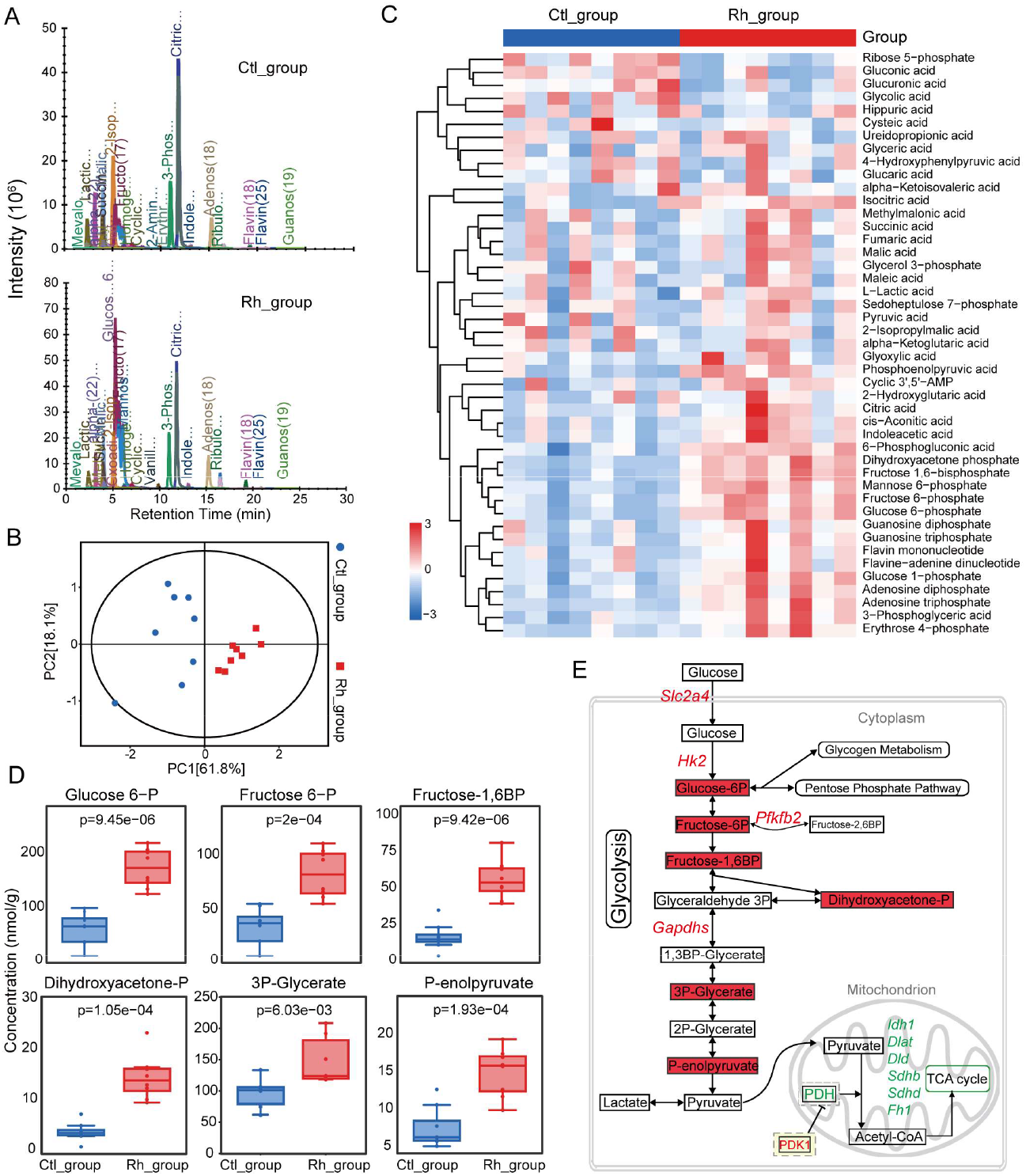
HPIC-MS/MS analysis of the central carbon metabolites in rhamnose-treated skin tissues. (A) Representative extracted ion chromatogram of metabolites in control or rhamnose-treated skin tissues. (B) PCA score plot of vehicle- and rhamnose-treated skin samples. (C) Heatmap of identified metabolites between control and rhamnose groups. (D) Quantitative analysis of significantly altered metabolites in glycolysis between control and rhamnose groups. (E) Scheme of glycometabolism pathway illustrating the changed metabolites, with the red color indicating highly upregulated genes and metabolites in the rhamnose-treated skin compared with the control, and the green color indicating downregulated ones. Ctl_group, Control group, Rh_group, Rhamnose group.

### 3.6 Rhamnose altered the transcriptional profile and induced metabolic reprogramming in hDPCs

To determine if rhamnose has similar effects on remodeling glycometabolism profiles in human hair follicles as it does in mice, we conducted a transcriptome analysis of hDPCs treated with 5 μM rhamnose for 24 hours. The results revealed that rhamnose significantly altered the gene expression profile of hDPCs (Figure 6A), with 175 upregulated genes and 248 downregulated genes. Gene ontology analysis showed that the upregulated genes were associated with several biological processes, including cellular response to hypoxia, gland development, tube morphogenesis, and positive regulation of cell migration (Figure 6B), while the downregulated terms were linked to the cellular response to oxygen and unfolded protein, as well as muscle and collagen development (Figure 6B). One of the key mechanisms that has evolved to cope with hypoxic conditions is the metabolic shift from predominantly relying on mitochondrial respiration to enhanced utilization of glycolysis in eukaryotic cells. This metabolic adaptation is crucial for sustaining sufficient ATP levels and ensuring cellular energy homeostasis (Kierans and Taylor, 2021). To investigate whether the upregulation of hypoxia response genes induced by rhamnose could alter glucose metabolism in hDPCs, we employed a Seahorse XFe24 analyzer to assess metabolic changes in the cells by monitoring the extracellular acidification rate (ECAR) and the oxygen consumption rate (OCR). The ECAR serves as a valuable parameter for assessing glycolytic activity through pH measurements that mainly reflect the production of lactate, an endpoint product of glycolysis, while OCR reflects the activity of mitochondria oxidative phosphorylation. As the results displayed (Figure 6C), rhamnose at a concentration of 5 μM enhanced key parameters of glycolytic functions, including glycolysis, glycolytic capacity, glycolytic reserve, and non-glycolytic acidification. Conversely, the OCR results showed that basal respiration, ATP production, maximal respiration, and spare respiration capacity were inhibited by rhamnose (Figure 6D). Overall, these findings demonstrated that rhamnose altered glycometabolism processes by increasing glycolytic capacity and attenuating mitochondrial respiration in hDPCs.

**Figure 6.**
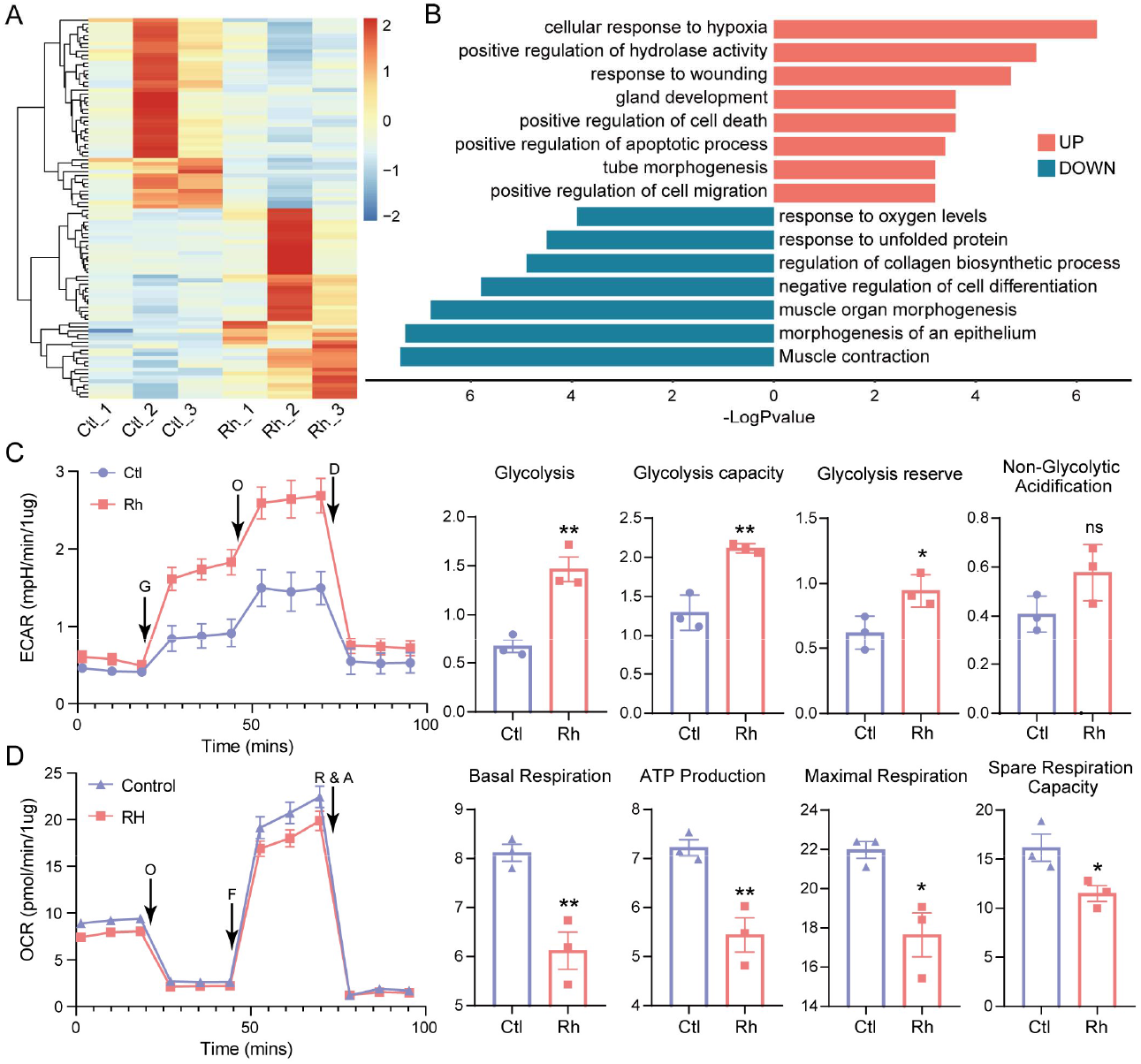
Transcriptional and metabolic analysis of hDPCs treated with rhamnose. (A) Heatmap analysis of the top 50 differentially expressed genes between the vehicle- and rhamnose-treated hDPCs. (B) GO enrichment analysis of up-and down-regulated genes in rhamnose-treated hDPCs. (C) Glycolytic stress tests for vehicle- and rhamnose-treated hDPCs (left), and the statistic analysis of key parameters: glycolysis, glycolytic capacity, glycolytic reserve, and non-glycolytic acidification. (D) Mitochondria stress tests for vehicle- and rhamnose-treated hDPCs (left), and statistic analysis of key parameters: basal respiration, ATP production, maximal respiration, and spare respiration capacity. Ctl, Control; Rh, Rhamnose. G, glucose; O, oligomycin; D, 2-deoxy-D-glucose; F, FCCP; R&A, rotenone/antimycin A. *P < 0.05, **P < 0.01.

## 4 Discussion

Small molecules have consistently driven medical breakthroughs and addressed unmet medical needs, as evidenced by recent advancements in the field. In the current study, we applied a novel approach by combining HTS^2^ and hair growth-inducing genes panel of hDPCs for high-throughput drug screening and identified rhamnose as a potential compound for promoting hair follicle growth. More importantly, we demonstrated the topical application of rhamnose accelerated hair follicle regeneration by stimulating the progression of the hair cycle to the anagen in mice. Furthermore, we found rhamnose changed the expression of molecules associated with glucose metabolism and induced a metabolic reprogramming favoring glycolysis in both mice skin and hDPCs. Rhamnose is a deoxy monosaccharide that is rarely found in animals but is commonly found in bacteria and plants in abundance, such as capsular polysaccharides, pectic polymers, and cell wall glycoproteins (Garcia-Vello et al., 2020, Jiang et al., 2021). Rhamnose has been shown to improve skin aging by thickening the epidermis, increasing procollagen I production, inhibiting the activity of elastase, hyaluronidase, and collagenase, as well as protecting skin fibroblasts from AGE-induced cytotoxicity (Novotná et al., 2023, Pageon et al., 2019, Ravelojaona et al., 2009). Our research broadens the scope of rhamnose applications in dermatology, investigates its potential as a therapeutic agent for hair loss, and provides a theoretical foundation for further drug development and clinical trials.

The development of specific drug screening models, efficient screening technologies, and comprehensive evaluation systems is crucial for enhancing drug screening efficiency. The restoration or enhancement of the hair follicle-inducing ability of hDPCs involves complex gene regulatory networks (Higgins et al., 2013), which makes it challenging to assess the effects of drugs on hDPCs through traditional single-gene detection methods. However, HTS^2^ technology overcomes the limitations of single gene assays by enabling the simultaneous quantitative detection of hundreds of genes closely associated with the hair follicle regenerative capacity of hDPCs. Moreover, due to limitations in sample availability and culture conditions, hDPCs cannot be extensively expanded *in vitro*. HTS^2^, with its ability to detect as little as 10 ng of total RNA and a minimal requirement of approximately 1000 cells, makes it possible to screen a larger number of drugs using limited cell samples. Additionally, HTS^2^ can be performed on in situ cell lysates with robotic automation, offering unparalleled stability and speed advantages compared to traditional methods. In conclusion, applying HTS^2^ technology to hDPCs models significantly enhances the efficiency of drug screening.

To explore the potential mechanism of rhamnose in modulating hair follicle growth, we believed that the alterations in gene expression profiles in rhamnose-treated skin after 14 days of treatment are more likely indicative of the premature anagen phenotype rather than a direct effect of rhamnose on skin cells. Consequently, we analyzed the *in vivo* transcriptome data from mouse skin treated with rhamnose for 7 or 10 days and *in vitro* data from rhamnose-treated hDPCs. Our analyses revealed DEGs induced by rhamnose are predominantly associated with hypoxia response and energy metabolism pathways. Under hypoxic conditions, eukaryotic cells have developed the ability to adapt to a metabolic switch to a heightened glycolytic rate, which drives significant alterations in cell behavior, phenotype, and fate (Hubbi and Semenza, 2015, Kierans et al., 2023, Macharia et al., 2021, Sathy et al., 2019). Hypoxia was reported to enhance the ability of DPCs to induce hair growth by triggering the generation of reactive oxygen species through the activity of nuclear NADPH oxidase 4 (Zheng et al., 2019). Previous works have suggested that robust glycolysis plays a vital role in hair follicle growth and regeneration. For instance, the expression of signature genes associated with hair induction and hair shaft elongation was decreased when glycolysis was inhibited, while lactate, an end product of glycolysis, showed the ability to activate HFSCs and induce hair growth (Choi et al., 2020, Flores et al., 2017, Sun et al., 2024). IM176OUT05, a refined biguanide, inhibits mitochondrial oxidative phosphorylation and enhances glycolysis, thereby promoting the hair follicle transition to the anagen phase and increasing hair follicle number in mice (Son et al., 2018). Here, we found rhamnose treatment enhanced glycolysis in hDPCs and mouse skin, which may represent one of the mechanisms through which rhamnose promotes hair growth.

As for the *in vivo* application of rhamnose, we acknowledge that rhamnose may exert an influence on the hair follicle epithelium, potentially contributing to its role in anagen induction. Future studies employing longitudinal single-cell transcriptomics and metabolic profiling of murine skin at various time points could elucidate the specific mechanisms underlying rhamnose’s promotion of hair growth. Furthermore, while the metabolic pathways of rhamnose have been extensively studied in microorganisms and plants, there is a paucity of research on its metabolism in mammals. The role of rhamnose as an intermediate metabolite in glycolysis within this context is yet to be elucidated, highlighting the need for additional research to determine its metabolic fate and its potential role in promoting hair growth in mammals.

## 5 Conclusion

In summary, our results suggest rhamnose promotes hair follicle growth and remodels glycometabolism both *in vivo* and *in vitro*. Our findings could be great news for those looking for alternative hair loss treatments, and further research could lead to new products that help manage this common condition.

## Author contributions

Conceptualization: ZY, DW, and RX; methodology: XL, YL, LL, SM, RC, and DW; software: XL, YL, and LL; investigation: ZY, XL, YL, LL, WJ, and YZ; writing—original draft: XL and ZY; writing—review and editing: ZY and RX; supervision: ZY, DW, and RX; funding acquisition: ZY and RX. All authors listed have made a substantial, direct, and intellectual contribution to the work and approved it for publication. All authors contributed to the article and approved the submitted version. The order of cofirst authors was determined by relative amount of data each contributed.

## References

1. Andl T, Zhou L, Zhang Y. The dermal papilla dilemma and potential breakthroughs in bioengineering hair follicles. Cell and tissue research 2023;391(2):221–33.

2. Chai M, Jiang M, Vergnes L, Fu X, de Barros SC, Doan NB, et al. Stimulation of Hair Growth by Small Molecules that Activate Autophagy. Cell reports 2019;27(12):3413-21.e3.

3. Choi M, Choi YM, Choi SY, An IS, Bae S, An S, et al. Glucose metabolism regulates expression of hair-inductive genes of dermal papilla spheres via histone acetylation. Scientific reports 2020;10(1):4887.

4. Driskell RR, Clavel C, Rendl M, Watt FM. Hair follicle dermal papilla cells at a glance. J Cell Sci 2011;124(Pt 8):1179–82.

5. Flores A, Schell J, Krall AS, Jelinek D, Miranda M, Grigorian M, et al. Lactate dehydrogenase activity drives hair follicle stem cell activation. Nature cell biology 2017;19(9):1017–26.

6. Gao Y, Jiang X, Wei Z, Long H, Lai W. The EDA/EDAR/NF-κB pathway in non-syndromic tooth agenesis: A genetic perspective. Frontiers in genetics 2023;14:1168538.

7. Garcia-Vello P, Sharma G, Speciale I, Molinaro A, Im SH, De Castro C. Structural features and immunological perception of the cell surface glycans of Lactobacillus plantarum: a novel rhamnose-rich polysaccharide and teichoic acids. Carbohydrate polymers 2020;233:115857.

8. Geyfman M, Plikus MV, Treffeisen E, Andersen B, Paus R. Resting no more: re-defining telogen, the maintenance stage of the hair growth cycle. Biological reviews of the Cambridge Philosophical Society 2015;90(4):1179–96.

9. Ginestet C. ggplot2: elegant graphics for data analysis. Oxford University Press; 2011.

10. Hao Y, Hao S, Andersen-Nissen E, Mauck WM, 3rd, Zheng S, Butler A, et al. Integrated analysis of multimodal single-cell data. Cell 2021;184(13):3573-87.e29.

11. Harel S, Higgins CA, Cerise JE, Dai Z, Chen JC, Clynes R, et al. Pharmacologic inhibition of JAK-STAT signaling promotes hair growth. Science advances 2015;1(9):e1500973.

12. Higgins CA, Chen JC, Cerise JE, Jahoda CA, Christiano AM. Microenvironmental reprogramming by three-dimensional culture enables dermal papilla cells to induce de novo human hair-follicle growth. Proceedings of the National Academy of Sciences of the United States of America 2013;110(49):19679–88.

13. Hubbi ME, Semenza GL. Regulation of cell proliferation by hypoxia-inducible factors. American journal of physiology Cell physiology 2015;309(12):C775–82.

14. Jiang N, Dillon FM, Silva A, Gomez-Cano L, Grotewold E. Rhamnose in plants - from biosynthesis to diverse functions. Plant science : an international journal of experimental plant biology 2021;302:110687.

15. Kierans SJ, Fagundes RR, Malkov MI, Sparkes R, Dillon ET, Smolenski A, et al. Hypoxia induces a glycolytic complex in intestinal epithelial cells independent of HIF-1-driven glycolytic gene expression. Proceedings of the National Academy of Sciences of the United States of America 2023;120(35):e2208117120.

16. Kierans SJ, Taylor CT. Regulation of glycolysis by the hypoxia-inducible factor (HIF): implications for cellular physiology. The Journal of physiology 2021;599(1):23–37.

17. Koralewicz MM, Szatkowska OA. Topical solutions for androgenetic alopecia: evaluating efficacy and safety. Forum Dermatologicum 2024.

18. Li H, Qiu J, Fu XD. RASL-seq for massively parallel and quantitative analysis of gene expression. Current protocols in molecular biology 2012;Chapter 4:Unit 4.13.1-9.

19. Love MI, Huber W, Anders S. Moderated estimation of fold change and dispersion for RNA-seq data with DESeq2. Genome biology 2014;15(12):550.

20. Macharia LW, Muriithi W, Heming CP, Nyaga DK, Aran V, Mureithi MW, et al. The genotypic and phenotypic impact of hypoxia microenvironment on glioblastoma cell lines. BMC cancer 2021;21(1):1248.

21. Madaan A, Verma R, Singh AT, Jaggi M. Review of Hair Follicle Dermal Papilla cells as in vitro screening model for hair growth. International journal of cosmetic science 2018;40(5):429–50.

22. McGinnis CS, Murrow LM, Gartner ZJ. DoubletFinder: Doublet Detection in Single-Cell RNA Sequencing Data Using Artificial Nearest Neighbors. Cell systems 2019;8(4):329-37.e4.

23. Novotná R, Škařupová D, Hanyk J, Ulrichová J, Křen V, Bojarová P, et al. Hesperidin, Hesperetin, Rutinose, and Rhamnose Act as Skin Anti-Aging Agents. Molecules (Basel, Switzerland) 2023;28(4).

24. Oh HA, Kwak J, Kim BJ, Jin HJ, Park WS, Choi SJ, et al. Migration Inhibitory Factor in Conditioned Medium from Human Umbilical Cord Blood-Derived Mesenchymal Stromal Cells Stimulates Hair Growth. Cells 2020;9(6).

25. Pageon H, Azouaoui A, Zucchi H, Ricois S, Tran C, Asselineau D. Potentially beneficial effects of rhamnose on skin ageing: an in vitro and in vivo study. International journal of cosmetic science 2019;41(3):213–20.

26. Ravelojaona V, Robert AM, Robert L. Expression of senescence-associated beta-galactosidase (SA-beta-Gal) by human skin fibroblasts, effect of advanced glycation end-products and fucose or rhamnose-rich polysaccharides. Archives of gerontology and geriatrics 2009;48(2):151–4.

27. Rendl M, Ban J, Mrass P, Mayer C, Lengauer B, Eckhart L, et al. Caspase-14 expression by epidermal keratinocytes is regulated by retinoids in a differentiation-associated manner. The Journal of investigative dermatology 2002;119(5):1150–5.

28. Sathy BN, Daly A, Gonzalez-Fernandez T, Olvera D, Cunniffe G, McCarthy HO, et al. Hypoxia mimicking hydrogels to regulate the fate of transplanted stem cells. Acta biomaterialia 2019;88:314–24.

29. Shim J, Park J, Abudureyimu G, Kim MH, Shim JS, Jang KT, et al. Comparative Spatial Transcriptomic and Single-Cell Analyses of Human Nail Units and Hair Follicles Show Transcriptional Similarities between the Onychodermis and Follicular Dermal Papilla. The Journal of investigative dermatology 2022;142(12):3146-57.e12.

30. Son MJ, Jeong JK, Kwon Y, Ryu JS, Mun SJ, Kim HJ, et al. A novel and safe small molecule enhances hair follicle regeneration by facilitating metabolic reprogramming. Experimental & molecular medicine 2018;50(12):1–15.

31. Sun D, Guan X, Moran AE, Wu LY, Qian DZ, Schedin P, et al. Identifying phenotype-associated subpopulations by integrating bulk and single-cell sequencing data. Nature biotechnology 2022;40(4):527–38.

32. Sun P, Wang Z, Li S, Yin J, Gan Y, Liu S, et al. Autophagy induces hair follicle stem cell activation and hair follicle regeneration by regulating glycolysis. Cell & bioscience 2024;14(1):6.

33. Wei G, Sun H, Wei H, Qin T, Yang Y, Xu X, et al. Detecting the Mechanism behind the Transition from Fixed Two-Dimensional Patterned Sika Deer (Cervus nippon) Dermal Papilla Cells to Three-Dimensional Pattern. International journal of molecular sciences 2021;22(9).

34. Wu T, Hu E, Xu S, Chen M, Guo P, Dai Z, et al. clusterProfiler 4.0: A universal enrichment tool for interpreting omics data. Innovation (Cambridge (Mass)) 2021;2(3):100141.

35. Yang K, Qiu T, Zhou J, Gong X, Zhang X, Lan Y, et al. Blockage of glycolysis by targeting PFKFB3 suppresses the development of infantile hemangioma. Journal of translational medicine 2023;21(1):85.

36. Zheng M, Jang Y, Choi N, Kim DY, Han TW, Yeo JH, et al. Hypoxia improves hair inductivity of dermal papilla cells via nuclear NADPH oxidase 4-mediated reactive oxygen species generation’. The British journal of dermatology 2019;181(3):523–34.

