## Supplemental Figure for "Rhamnose-mediated modulation of hair follicle growth: insights from chemical genomics"

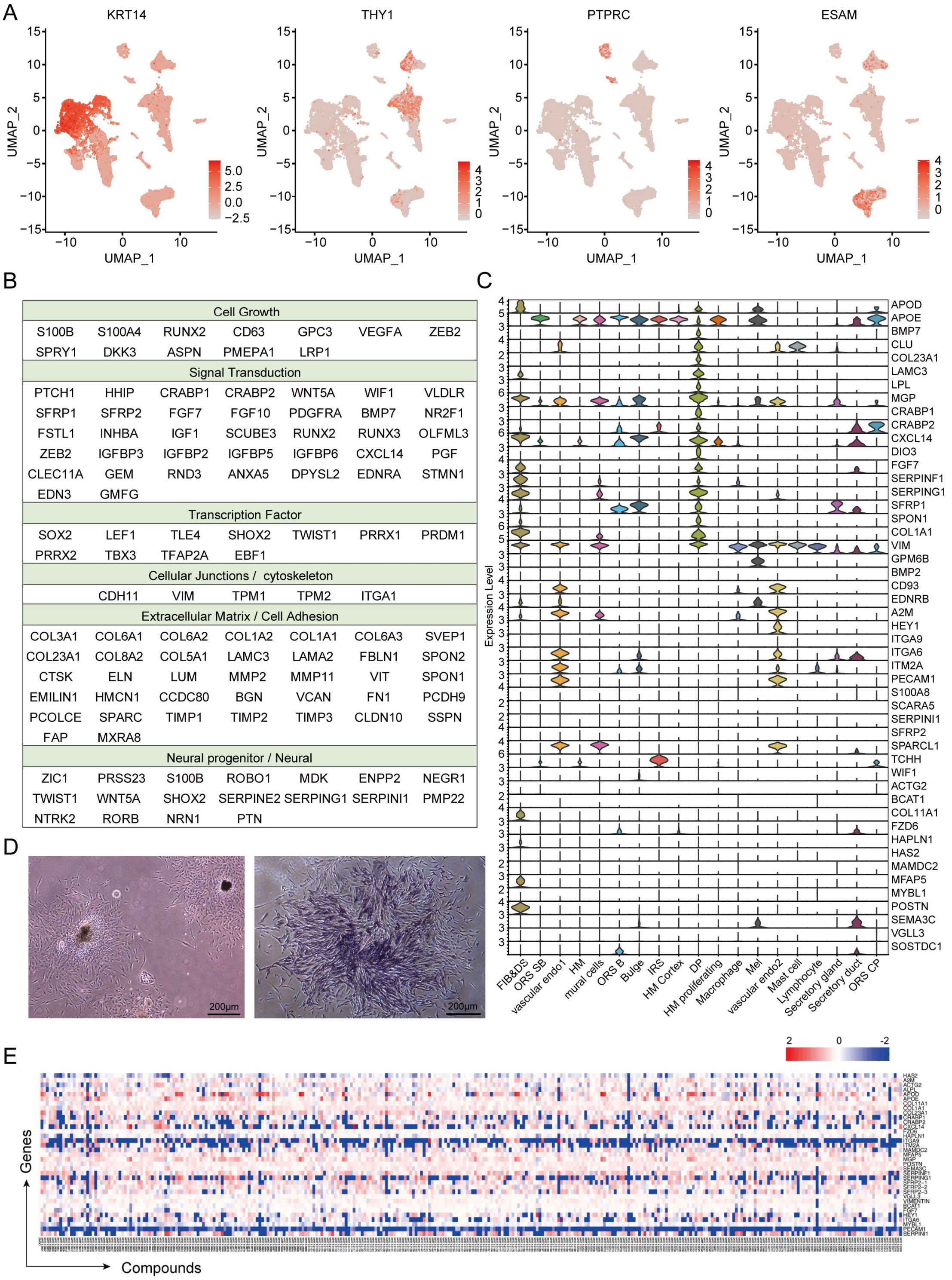


**Figure S1 Single-cell data analysis of human scalp tissue and HTS$^2$ screening result**

(A) Feature plots showing the expressions of selected cell type-specific genes. (B) Signature genes identified in the DP cluster. (C) The violin plots showing the expressions of panel genes for HTS$^2$ screening in the single-cell dataset. (D) The morphology of freshly isolated and adherently cultured dermal papilla for 9 days (left) and ALP staining of cultured hDPCs (right). (E) Heatmap displaying the expression profiles of hair follicle induction-related genes perturbed by thousands of small molecular compounds. Ave.Exp, Average expression; Pct.exp, Percent expressed.

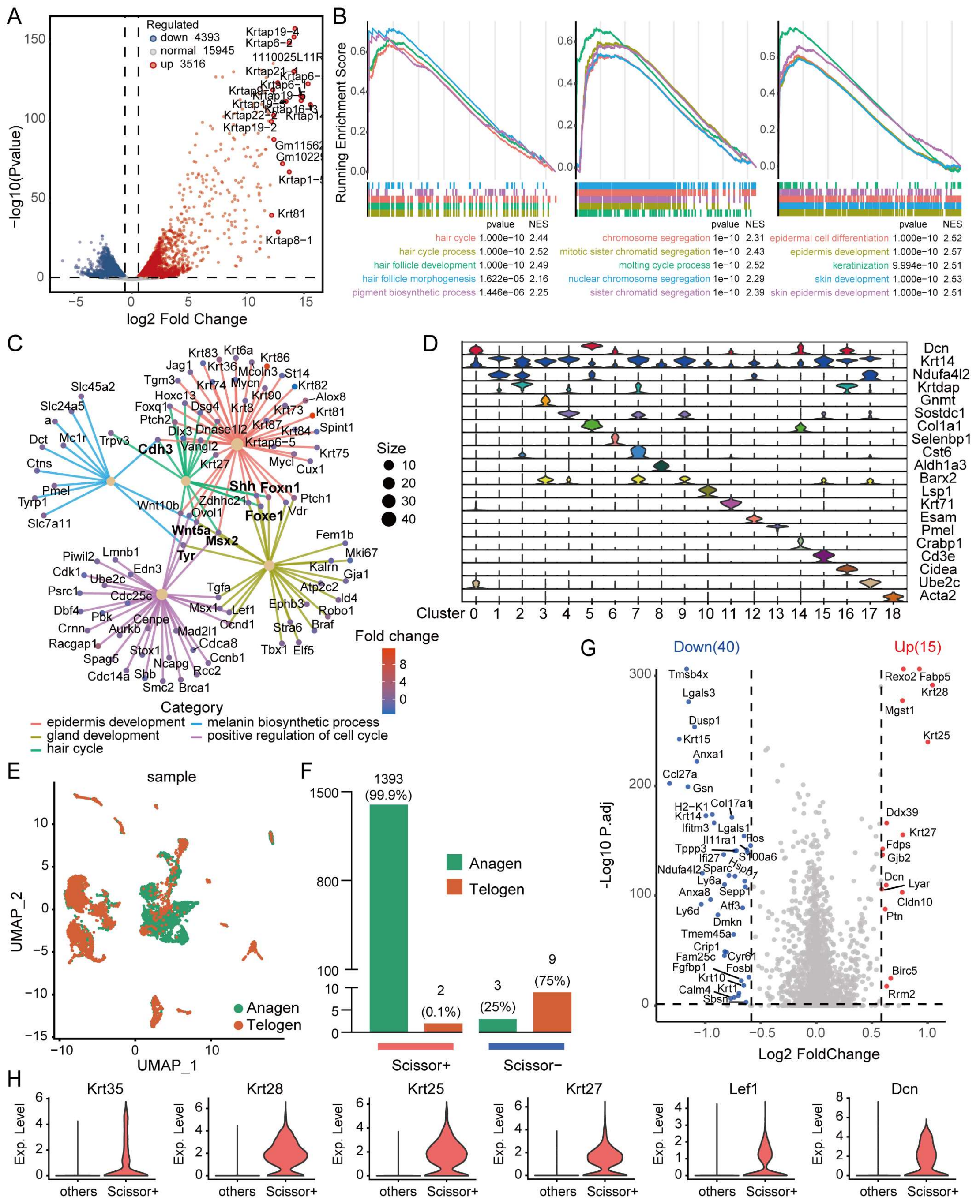


**Figure S2 Identification of cell subpopulations and molecule changes in mice skin after rhamnose-treated 14 days**

(A) Volcano plot showing DEGs in the rhamnose-treated group relative to the pre- and vehicle-treated groups at 14 days. (B) GSEA plots of the highly enriched genes in the rhamnose-treated group at 14 days. (C) CNET plot of the representative biological processes from rhamnose-treated 14-day mice. (D) Violin plots depicting the expression patterns of marker genes across each identified cluster. (E) The UMAP clustering of cell populations from different samples. (F) Bar plots showing the detailed phenotypic constitutions of the scissor-selected cells. (G) The volcano plot of DEGs in Scissor+ cells versus all other cells. DEGs are genes with adjusted p-values < 0.05 and absolute Fold Change ⩾1.5. (H) The violin plots illustrating the significantly upregulated genes in Scissor+ cells.

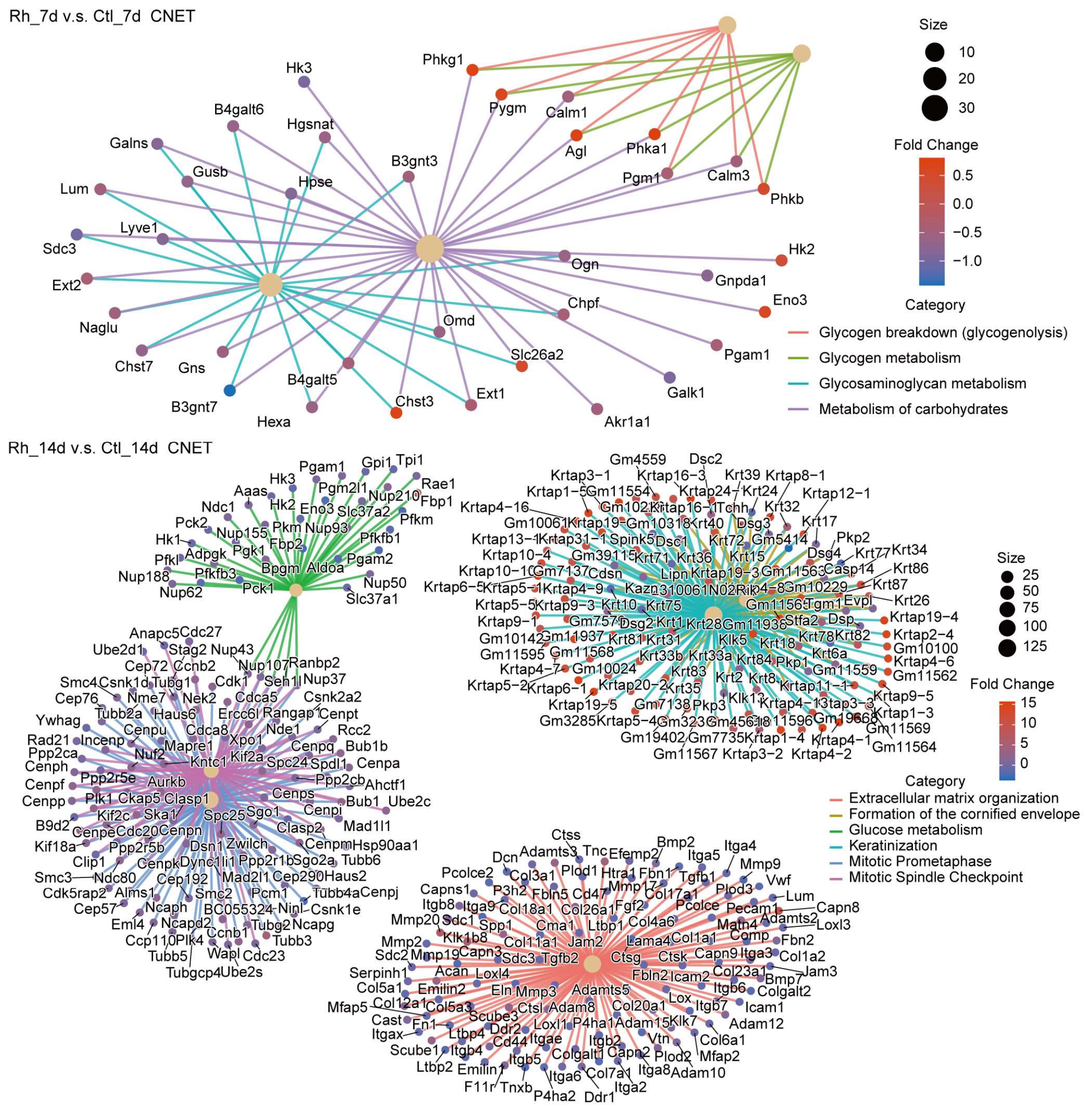


**Figure S3** CNET plots illustrating the key Reactome pathways in the rhamnose-treated groups at 7 days (top) and 14 days (bottom), in comparison to the control.

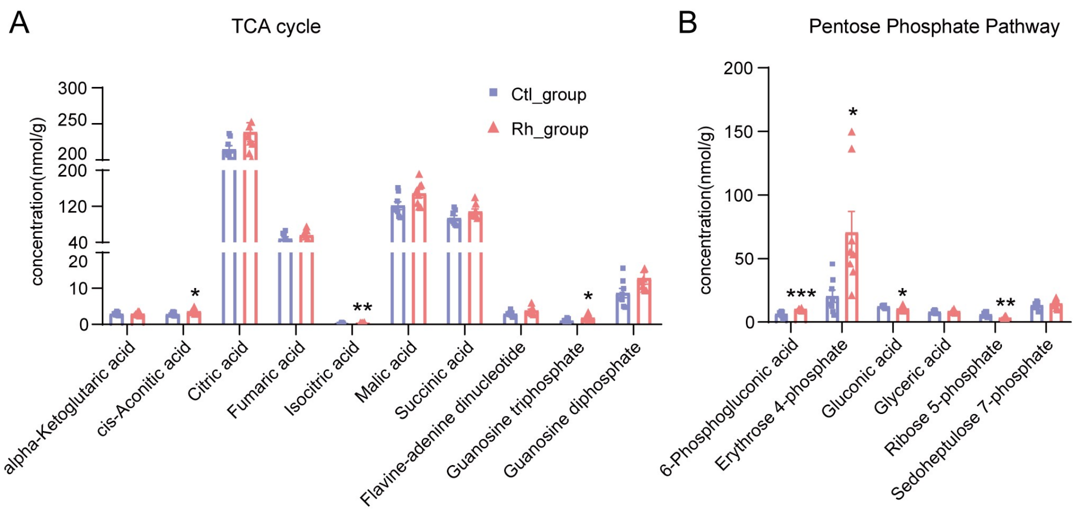


**Figure S4 Quantification of metabolites associated with the TCA cycle and pentose phosphate pathway.** Bar plots showing the concentrations of metabolites in TCA cycle (A) and pentose phosphate pathway (B) between vehicle- and rhamnose-treated group. *P < 0.05, **P < 0.01, ***P < 0.001. Ctl_group, Control_group, Rh_group, Rhamnose_group.
